# Genomic patterns of divergence in the early and late steps of speciation of the deep-sea vent thermophilic worms of the genus *Alvinella*

**DOI:** 10.1101/2022.02.28.482349

**Authors:** Camille Thomas–Bulle, Denis Bertrand, Niranjan Nagarajan, Richard Copley, Erwan Corre, Stéphane Hourdez, Éric Bonnivard, Adam Claridge-Chang, Didier Jollivet

## Abstract

**Background:** The transient and fragmented nature of the deep-sea hydrothermal environment made of ridge subduction, plate collision and the emergence of new rifts is currently acting to the separation of vent populations, local adaptation and is likely to promote bursts of speciation and species specialization. The tube-dwelling worms *Alvinella pompejana* called the Pompeii worm and its sister species *A. caudata* live syntopically on the hottest part of deep-sea hydrothermal chimneys along the East Pacific Rise. They are exposed to extreme thermal and chemical gradients, which vary greatly in space and time, and thus represent ideal candidates for understanding the evolutionary mechanisms at play in the vent fauna evolution.

**Results:** In the present study, we explored genomic patterns of divergence in the early and late steps of speciation of these emblematic worms using transcriptome assemblies and the first draft genome to better understand the relative role of geographic isolation and habitat preference in their genome evolution. Analyses were conducted on allopatric populations of *Alvinella pompejana* (early stage of separation) and between *A. pompejana* and its syntopic species *Alvinella caudata* (late stage of speciation). We first identified divergent genomic regions and targets of selection as well as their position in the genome over collections of orthologous genes and, then, described the speciation dynamics by documenting the annotation of the most divergent and/or positively selected genes involved in the isolation process. Gene mapping clearly indicated that divergent genes associated with the early stage of speciation, although accounting for nearly 30% of genes, are highly scattered in the genome without any island of divergence and not involved in gamete recognition or mito-nuclear incompatibilities. By contrast, genomes of *A. pompejana* and *A. caudata* are clearly separated with nearly all genes (96%) exhibiting high divergence. This congealing effect however seems to be linked to habitat specialization and still allows positive selection on genes involved in gamete recognition, as a possible long-duration process of species reinforcement.

**Conclusion:** Our analyses pointed out the non-negligible role of natural selection on both the early and late stages of speciation in the emblematic thermophilic worms living on the walls of deep-sea hydrothermal chimneys. They shed ligth on the evolution of gene divergence during the process of speciation and species specialization over a very long period of time.

## Background

The transient and fragmented nature of the deep-sea hydrothermal environment made of ridge subduction, plate collision and the emergence of new rift systems likely led to the separation of vent communities at a large spatial scale with transient events of spatial isolation and bursts of speciation [1–4]. Most species inhabiting these unstable and harsh environments often display a rapid growth, a fast maturation, a high investment in reproduction relative to survival and good dispersal abilities enabling the colonization of vent emissions as soon as they appear [5–6]. Many studies notably focused on how this highly specialized and endemic fauna disperse and colonize new territories in the face of habitat fragmentation [7–8]. Given the dynamics of the vents associated with the tectonic activity along oceanic ridges and its effect on the metapopulation dynamics [9], such environment appears ideal to study the mechanisms by which speciation occurs and to tease apart the relative role of geography and local adaptation to this extent. On one hand, the vent environment is highly variable with abrupt chemical gradients leading to a mosaic of habitats where species are spatially and temporally partitioned [10–12]. This may likely promote ecological speciation when migration is not able to counter-balance local selection at some specific locations. On the other hand, populations are connected within a linear network of habitats and tectonic plates rearrangements are likely to result in either species spatial isolation or secondary contact zones where hybridization can occur [3, 13–16]. In addition to isolation by distance between vent populations, patterns of genetic differentiation are mainly explained by the presence of physical barriers to dispersal such as transform faults, microplates, triple junction points or zones of weaker hydrothermal activity [3,15–16]. These relatively dynamic geological features may produce vicariant events in the faunistic composition of vent communities [17] that often coincide with the appearance of barriers to gene flow and hybrid zones [14,16,18].

Early incipient species often represent groups of individuals that can still interbreed and exchange genes despite the emergence of pre- or post-zygotic barriers to gene flow. During the process of speciation and even at the latest steps of reinforcement, emerging species are thus not a collection of impermeable taxonomic units with distinct morphologies but rather dynamic entities gradually heightening their reproductive barriers in a more or less continuous way over long periods of time [19]. This process depends on the divergence history of the populations and their associated ecological changes. But it is worth mentioning that exceptions where speciation happens relatively fast also exist, for example in the case of genomes under disruptive selective conditions following changes in ploidy [20–21] or rapid shifts in reproductive timing/behaviour due to the colonization of new territories/habitats [22–23]. In general, pre- and post-zygotic mechanisms leading to reproductive isolation result in a gradual accumulation of genetic barriers during the separation of groups of individuals in either space or time [24–31]. The point at which speciation is initiated and then considered complete is however vague and varies with the species concept used (*e.g.*, complete reproductive isolation, hybrid counter-selection, phylogenetic isolation with complete sorting of allelic lineages between populations). In the initial part of the speciation process, individuals are likely to still exchange genes across semi-permeable barriers resulting in heterogeneous migration rates along the genome but could also endure local adaptation resulting in a loss of genetic diversity over some portions of their interacting genomes [32].

Recent studies on the genetic architecture of reproductive isolation across the entire genome often found a genome-wide heterogeneity of genetic differentiation between populations, ecotypes or even well-separated morphological species, with highly divergent regions of the genome contrasting with regions of lower divergence rather than a homogeneous rate of differentiation across the genome [30, 33–34]. To this extent, genome scans of differentiation and genome-wide linkage maps based on polymorphic sites at multiple loci across the genome, represent powerful tools to address the genomic landscape of population/species isolation. Regions displaying high levels of differentiation between lineages, so-called “genomic islands of speciation” were logically assumed to exhibit loci underlying reproductive isolation when the time elapsed since isolation was long enough to create genetic incompatibilities [35]. Most of the time, “islands of speciation” are believed to result from local barriers to gene flow at some specific genes that rapidly led to allele fixation, gene hitchhiking and the subsequent accumulation of divergence [36]. These islands differ from incidental islands of divergence resulting from accelerated rates of lineage sorting within populations due to recurrent events of either selective sweeps or background selection not necessarily related to reproductive isolation [37]. The genomic island metaphor has proven popular and a wide array of studies searching for “islands of speciation” in multiple taxa has been published over the last decade [30,34,35,38–42]. These studies identified outlier loci or regions that stand out from the distribution of the multi-loci genetic differentiation (most often through F_ST_ scans) expected under models of migration with or without isolation and heterogeneous rates of migration across genomes (*e.g.*, assortative mating, hybrid counter-selection, divergent selection). Such outlier genes usually fall into three categories: (1) outliers resulting from a differential effect of purifying selection in non-crossover regions of the separating genomes, (2) outliers involved in Dobzhansky-Muller incompatibilities that affect the fitness of hybrids/heterozygotes, and (3) outliers resulting from differential local adaptation between the splitting lineages. In the last category, a limited number of alleles involved in local adaptation have been found to carry a selective value strong enough to initiate a barrier to gene flow primarily by divergent selection. When considering speciation from both ends of the road, how do these patterns of divergence differ between the early and the late stages of isolation?

To better understand the role of genomic architecture on speciation, the number, width, and level of clustering of divergent regions within the genome of early incipient species are usually compared with more divergent congeneric pairs. It is expected that early in speciation, regions of divergence should be small and randomly scattered in the genome and possibly restricted to a few non-recombining regions. These regions may correspond to the accumulation of genetic incompatibilities or to loci under diversifying selection strong enough to overcome gene flow which would be expected to homogenize the rest of the genome. Conversely, later in speciation, when gene flow is greatly reduced between the two genomes by the cumulated effect of multiple genetic barriers, divergent selection, physical linkage between targets of selection and chromosomal rearrangements may create longer genomic regions of divergence. Such processes acting over larger regions of the genome, followed by gene specialization within each genome in response to local adaptation contribute to a genome-wide “congealing” (GWC) effect that finalizes speciation with reproductive isolation [43–46]. GWC refers to the period along the speciation continuum when the genomes of the splitting populations essentially become sealed and when reproductive isolation becomes a property of the whole genome not just of a few loci [43]. Therefore, incomplete lineage sorting and semi-permeable barriers to gene flow (associated with the early stages of speciation) and the wide synergetic effects of genome congealing due to chromosomal linkage between diverging genes (associated with the latest stages) are likely to result in very distinct patterns of divergence at a given stage of the speciation process. Moreover, positive selection over the genome due to gene specialization under local conditions should affect KEGG pathways differently over the course of the splitting process. Although fundamental to our understanding of speciation, such comparisons between the early and late stages of speciation, and the time of transition between these two stages have rarely been done when looking at the relationship between the number of diverging genes in association (congealing effect) and their accumulated divergence (time since species separation).

hydrothermal chimneys along the East Pacific Rise (EPR) (Figure 1) [47–48]. Using transcriptome assemblies, we carefully examined the relative role of geographic isolation and habitat preference in shaping gene divergence between newly isolated populations (in allopatry) of *Alvinella pompejana* and between this latter species and its syntopic species *Alvinella caudata* (niche specialization & reinforcement in sympatry). The tube-dwelling worms *Alvinella pompejana* Desbruyères & Laubier 1980 [49] (Figure 1C and 1D) called the Pompeii worm and its sister species *A. caudata* Desbruyères & Laubier 1986 [50] (Figure 1B) live on the wall of still-hot chimneys of the EPR from 23°N (Guaymas basin) to 38°S (Pacific-Antarctic Ridge). The first description of these two worms considered them as two morphologically distinct ontogenic forms of a single species, one juvenile form with a long tail bearing filamentous campylobacteria on modified notopodia (corresponding to *A. caudata*) and the adult reproductive form without tail but bearing the same epibionts on the whole dorsal part of its body (corresponding to *A. pompejana*). But enzyme polymorphism analyses led to the subdivision of this taxon into two distinct species without any shared electromorphs between them [51–52]. For both species, a strong physical barrier to gene flow was depicted at the Equator near 7°S/EPR and separates Northern and Southern vent populations in two allopatric groups of putative cryptic species with differentiation particularly marked in the Pompeii worm *Alvinella pompejana* [3, 15]. Differences in the vent-site turn-over from both sides of the EPR and its subsequent effect on local hydrothermal vent conditions (*i.e.* changes in thermal conditions due to a highest proportion of still-hot chimneys and fluid chemistry heterogeneity at both local and regional scales) could represent favorable conditions for local adaptation and subsequent ecological filtering when migration is low or episodic across the barrier. In addition, depth gradient along the ridge or between ridges, extreme variations in the hydrothermal fluid composition and subsequent effects on symbiotic associations with divergent bacterial strains are likely to induce divergent selection between vent populations [53].

**Figure 1:**
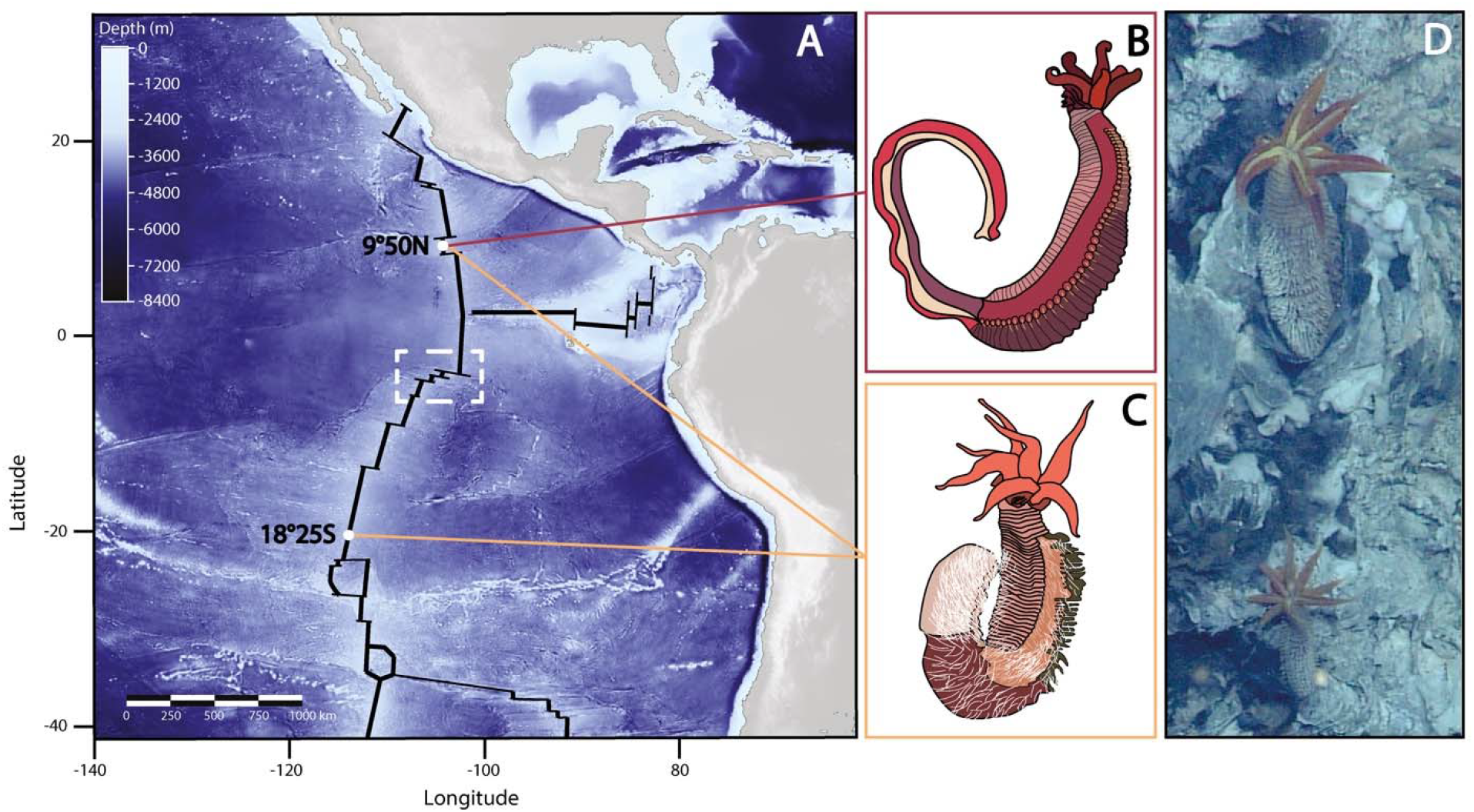
A) Bathymetric map of the East Pacific Ridge (in black), sampled populations sites along the ridge (white dots) and the geographical barrier between Southern and Northern populations (white dashed square); B) Drawing of a specimen of *Alvinella caudata;* C) Drawing of a specimen of *Alvinella pompejana*; D) *A. pompejana*, populations in situ (East Pacific Rise: 13°N, 2630 m); PHARE cruise

To compare both ends of the speciation process, we performed the first genome scans of both synonymous and non-synonymous gene divergences using transcriptome assemblies of the two allopatric forms of *A. pompejana* that have recently separated about 1.5 Mya (early stage of speciation), and that of its co-occurring sister species *A. caudata,* known to exhibit almost complete reproductive isolation with *A. pompejana* (late stage of speciation). We then identified divergent genomic regions and targets of selection as well as their position in the genome over collections of orthologous genes and, thus, described the speciation dynamics by documenting the annotation of the most divergent and/or positively selected genes involved in the species isolation.

## Results

### Assembly and annotation of the reference genome and associated transcriptomes

The assembled genome of *A. pompejana* consisted in 3,044 scaffolds for a total size of 245Mb, which represents 61% of the total genome size (400 Mb). The mapping of both divergence and associated d_N_/d_S_ along scaffolds was however only performed on the 113 longest scaffolds (> 300,000 bp) to maximize the number of genes per scaffold.

### Identification of orthologous genes

Using the genome of *A. pompejana* as a reference database, we analyzed 94,006 pairwise Blastn alignments between the Northern and Southern transcriptomes of *A. pompejana* and 68,963 pairwise Blastn alignments between the transcriptomes of *A. pompejana* and *A. caudata*. As these putative genes contain paralogs, tandemly repeated paralogs, allelic forms of a gene in either genes with or without introns, we filtered our datasets to only keep a subset of the most likely orthologous genes. This reduces the number of pairwise alignments to 22,558 and 48,134 respectively. Determination and translation of the CDS with a minimum threshold length of 300 bp and sequence-similarity search against the Uniprot database reduced the two final datasets to 6,916 and 6,687 pairwise alignments of orthologous coding sequences on which further analyses were performed.

### Distribution of genetic divergence and d_N_/d_S_ values

The evolutionary dynamics of genes during speciation was investigated by calculating pairwise synonymous to non-synonymous substitution rates and the associated divergence between orthologous transcripts of divergent individuals within and between closely-related species (see the “Methods” section). Among the 6,916 orthologous genes studied between the two Northern and Southern individuals of *A. pompejana*, 58% of the coding sequences are strictly identical with no nucleotide differences, 26% of the genes display divergences between 0.5% and 1.75% (0.01>t>0.05), and only 4% of the genes represent outliers that diverge by more than 2% (t>0.05) along their sequence (Figure 2). As expected in early speciation steps, the overall mean divergence of genes is low (t=0.016) with values following a negative exponential distribution with a long tail of outliers. However, the distribution has a slight bimodal shape, the second unexpected mode being centered around a value ten times greater (i.e., about 0.020 substitution per codon) (Figure 2). This bimodal shape was not encountered when assessing the intra-specific allelic diversity between two Northern individuals (average diversity = 0.0096 ± 0,0051 with fewer outliers (t>0.05 =2%), see Figure 2). Analysis of both transition and transversion rates between individuals from North and South were 4 times greater than those found by comparing two distinct individuals of the same population in the northern part of the EPR (Figure 3).

**Figure 2:**
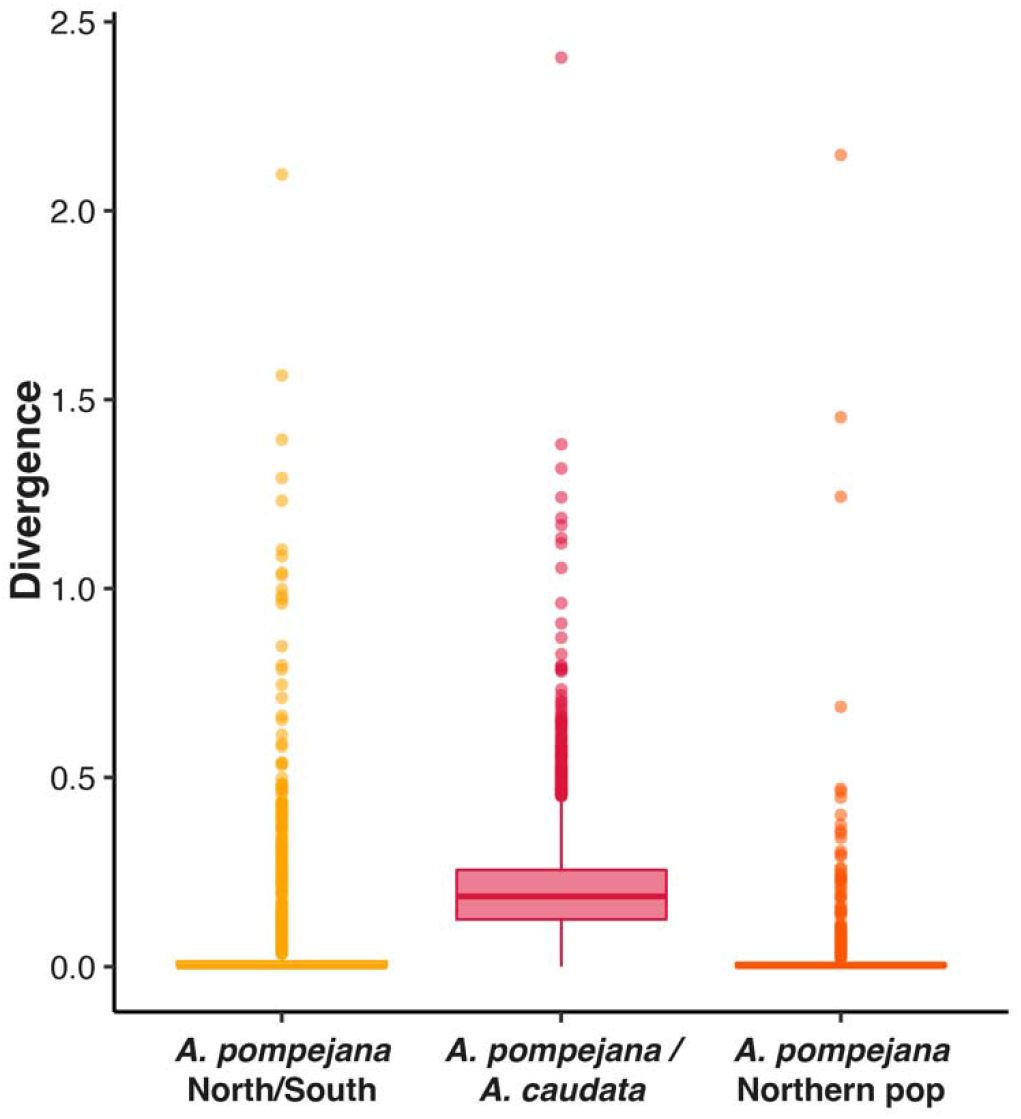
Boxplots of the distribution of the divergence values found in different stages of speciation in *Alvinella spp*. worms. The yellow color corresponds to the comparison *A. pompejana south/north*. The red color corresponds to the comparison *A. caudata/A. pompejana.* The orange color corresponds to the comparison *A. pompejana* Northern populations.

**Figure 3:**
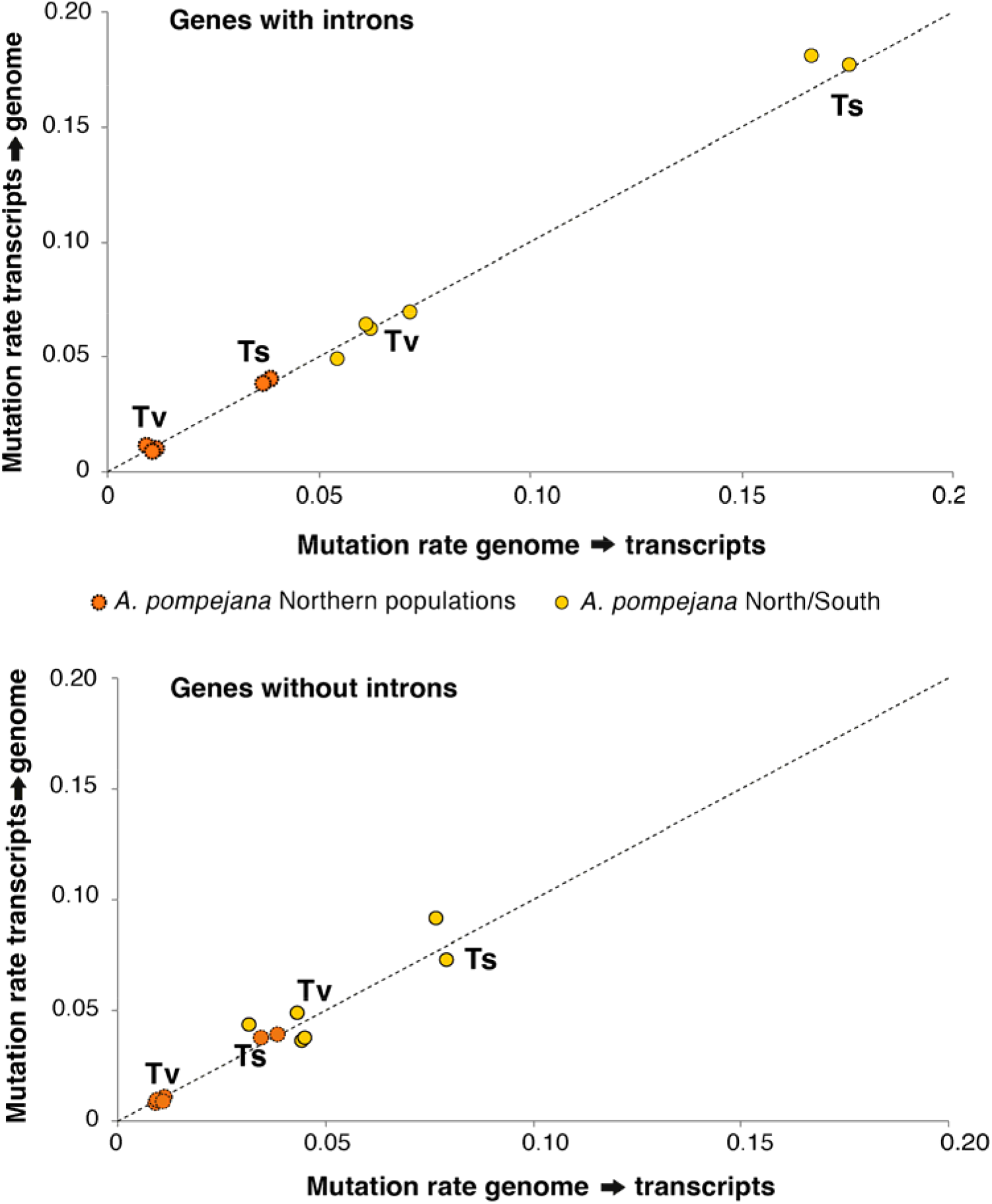
Transition and translation rates obtained between orthologous pairs of genes of *Alvinella pompejana* by comparing single-individual transcriptomes from the north (9°50N) and south (18°25S) EPR onto the genome of the worm (northern individual). Rates were obtained from coding sequences obtained from the orthologous gene datasets used for the d_N_/d_S_ analysis by discriminating genes without introns and genes with introns (i.e. the exonic regions of genes obtained after mapping transcripts onto the genome).

On the opposite, the large majority of the 6,687 genes are highly divergent between *A. pompejana* and *A. caudata*. Divergence follows a Gaussian distribution centered around 0.2 substitution per codon with an asymmetric tail of outliers (Figure 2). Only 0.4% of the sequences are strictly identical with no nucleotide differences corresponding to only 33 pairwise alignments, 3% of the genes display divergence levels between 10% and 25% (0.25>t>0.75), and 96% of the genes diverge by more than 25% (t>0.75) along their sequence. Thus, the distribution of the divergence between *A.caudata* and the 2 populations of *A.pompejana* is significantly different (Figure 2; Wilcoxon test, p < 2.2e-16), but this distribution is not significantly different within *A. pompejana* populations (Wilcoxon test, p = 0.8266).

Regarding the distribution of the d_N_/d_S_ values between North and South individuals of *A. pompejana*, most of the genes are non-divergent with a d_N_/d_S_ ratio set to zero (Figure 4). Among diverging genes, 44% of the genes are under strong purifying selection with d_N_/d_S_ values lower than 0.25. 36% of the d_N_/d_S_ values are distributed in a small peak between 0.25 and 0.5, and 6% of the genes are under positive selection with d_N_/d_S_ values higher than 1 and represent about 3% of the total number of genes examined (Figure 4). This peak of d_N_/d_S_ was not encountered when comparing the two northern *A. pompejana* transcriptomes although the number of outliers (d_N_/d_S_>1) was higher between sequences of these two specimens (about 12%). The absolute values of d_N_/d_S_ and π_N_/π_S_ were greater than the 2% of genes under positive selection (d_N_/d_S_>1) found between *A. pompejana* and *A. caudata*, probably because of a sampling bias in estimating both synonymous and non-synonymous rates from poorly intra-specific divergent sequences. The remaining d_N_/d_S_ values lesser than one between *A. pompejana* and *A. caudata* (98%) contains about 30% of genes with d_N_ equal to zero (*i.e.*, “frozen” proteins), about 53% of genes with d_N_/d_S_ values ranging from 0 to 0.25 (genes under strong to moderate purifying selection) and 15% of genes evolving under more relaxed conditions (close to one). The distribution of d_N_/d_S_ values within and between species was significantly different (Wilcoxon test, p < 2.2e-16) with a clear bi- to tri-modal distribution of the d_N_/d_S_ values in the specific case of the within-species comparison. The average d_N_/d_S_ estimated for both early and late steps of speciation was nearly equal with values of 0.14 and 0.17, respectively. In other words, at least 86% and 84% of non-synonymous mutations are deleterious and do not fix in the two *Alvinella* worms. This represents a minimum estimate of the fraction of strongly deleterious mutations because even a small number of advantageous mutations will contribute disproportionately to divergence [54].

**Figure 4:**
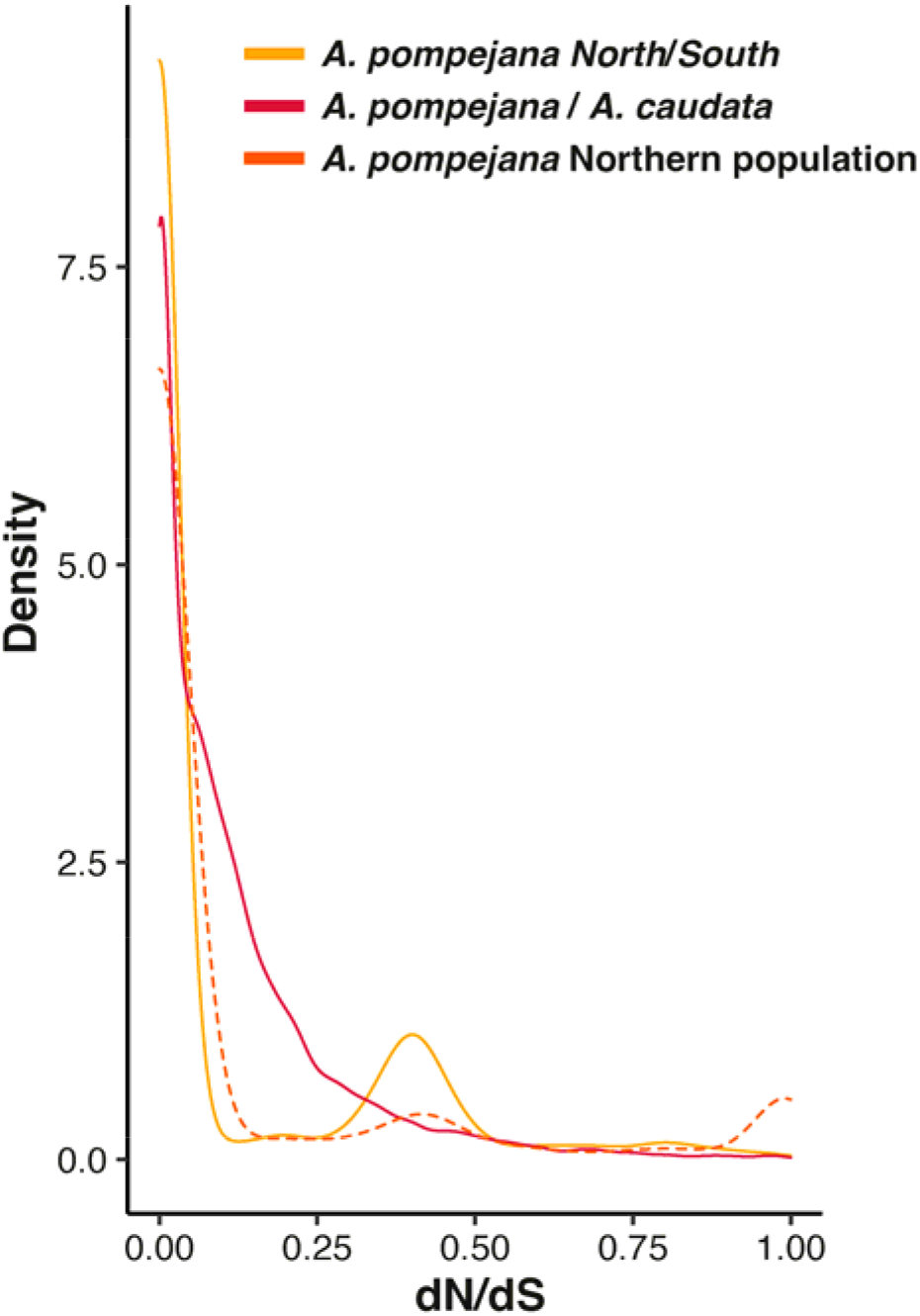
Density distributions of d_N_/d_S_ values estimated for each pairwise alignment of orthologous genes in both the early and late states of speciation in *Alvinella spp*. worms using the method of Nielsen & Yang (1998) implemented in PamlX. The yellow color corresponds to the comparison *A. pompejana south/north*. The red color corresponds to the comparison *A. caudata/A. pompejana.* The orange color corresponds to the comparison *A. pompejana* Northern populations.

### Functions of positively selected genes and genes with very high evolutionary rates

#### Early stage of speciation (A. pompejana south/north comparison)

Among the 41 genes under positive selection (d_N_/d_S_ >1) for which d_N_/d_S_ values remained significant after the resampling test of the d_N_/d_S_ difference and the FDR procedure, only 12 of them were annotated in the UniProt database. These genes are encoding proteins involved in (1) transcription/translation/replication regulation and biosynthesis (ribosomal protein rpl19, GTP-binding protein 1, elongation factor 1alpha, zinc finger protein GLI1) some of which being involved in a wide variety of biological functions including spermatogenesis, (2) endocytosis and immunity regulation, which may play a crucial role in the worm’s epibiosis (Rab5 protein, CD209 antigen protein A), (3) neuro-transmission regulation (complexin) and the development of sensory organs (protein mab-21), (4) development of nephridies, and possibly gonoducts in alvinellids (actin-binding Rho protein), (5) mRNA methylation (pre-miRNA 5’-monophosphate methyltransferase), and (6) setae/modified hook formation that should influence reproductive/thermoregulatory behavior (chitin synthase).

#### Late stage of speciation (the A. pompejana/A. caudata comparison)

In the same way, among the 102 orthologous genes between *A. caudata* and *A. pompejana* for which d_N_/d_S_ values were greater than one, only 37 remained significant after the resampling test of the d_N_/d_S_ difference and the FDR procedure but, in this case, most them were annotated (31 genes) in the UniProt database. These genes encode for proteins involved in (1) carbohydrate catabolism (deoxyribose-phosphate aldolase), (2) immunity (innate and adaptive), viral responses, apoptosis (fibrinogen-like protein, lysosomal protective protein, acetylcholine receptor), (3) steroid/lipid metabolism and membrane composition (transmembrane protein, 3-oxo-5-alpha-steroid 4-dehydrogenase, methylsterol monooxygenase, protein DD3-3-like), and tegument structure (actin, collagen, epidermal growth factor named fibropellin, GTP-binding protein, protein-methionine sulfoxide oxidase MICAL1), (4) sexual differentiation, spermatogenesis/oogenesis, sperm adhesion (zonadhesin, innexin), (5) DNA damage repairs and methylation (ATP-dependent RNA helicase DDX51 and DDX58, poly-(ADP-ribose) polymerase, forkhead box G1 protein WD repeat-containing protein), oxidative stress response and protein glycosylation (beta-1.3-galactosyltransferase 5-like, C19orf12), (6) cell proliferation/apoptosis, mitosis, neuron formation (Tumor Necrosis Factor alpha, ras-related protein, APC/C activator Cdc20 protein, tyrosine-protein kinase, protein FAM134C, zinc finger protein), and (7) signaling mediated by metals (metabotropic glutamate receptor). Despite a suspected several millions of years of species separation, it is worth-noting that positive selection is still acting on genes involved in reproductive isolation, namely on zonadhesin and innexin, hence possibly reinforcing species boundaries.

### Patterns of nucleotide substitutions and highly divergent genes

Because natural selection usually operates mostly on the non-synonymous sites (but also on translational accuracy [55]), the relative values of d_N_ and d_S_ may provide a key for making inferences about the adaptive evolution of protein-coding genes. Usually, the synonymous rate is three times greater than the rate of substitutions at nonsynonymous sites but should be equal under neutral evolution. Thus, the depression in non-synonymous substitution rate is interpreted as being caused by natural selection eliminating deleterious mutations to comply with both the 3D structure and functions of proteins. The relationship between these two relative rates of substitutions was assessed in the early and late steps of speciation thanks to a linear regression to take into account the very high variance of these rates. The two comparisons greatly differ in the slope and intercept of the linear regression between d_N_ and d_S_ (*A. pompejana* North/South: y= 0.002631+ 0.040525x; *A. pompejana / A. caudata*: y = 0.01818 + 0.01186x) (Figure 5).

**Figure 5:**
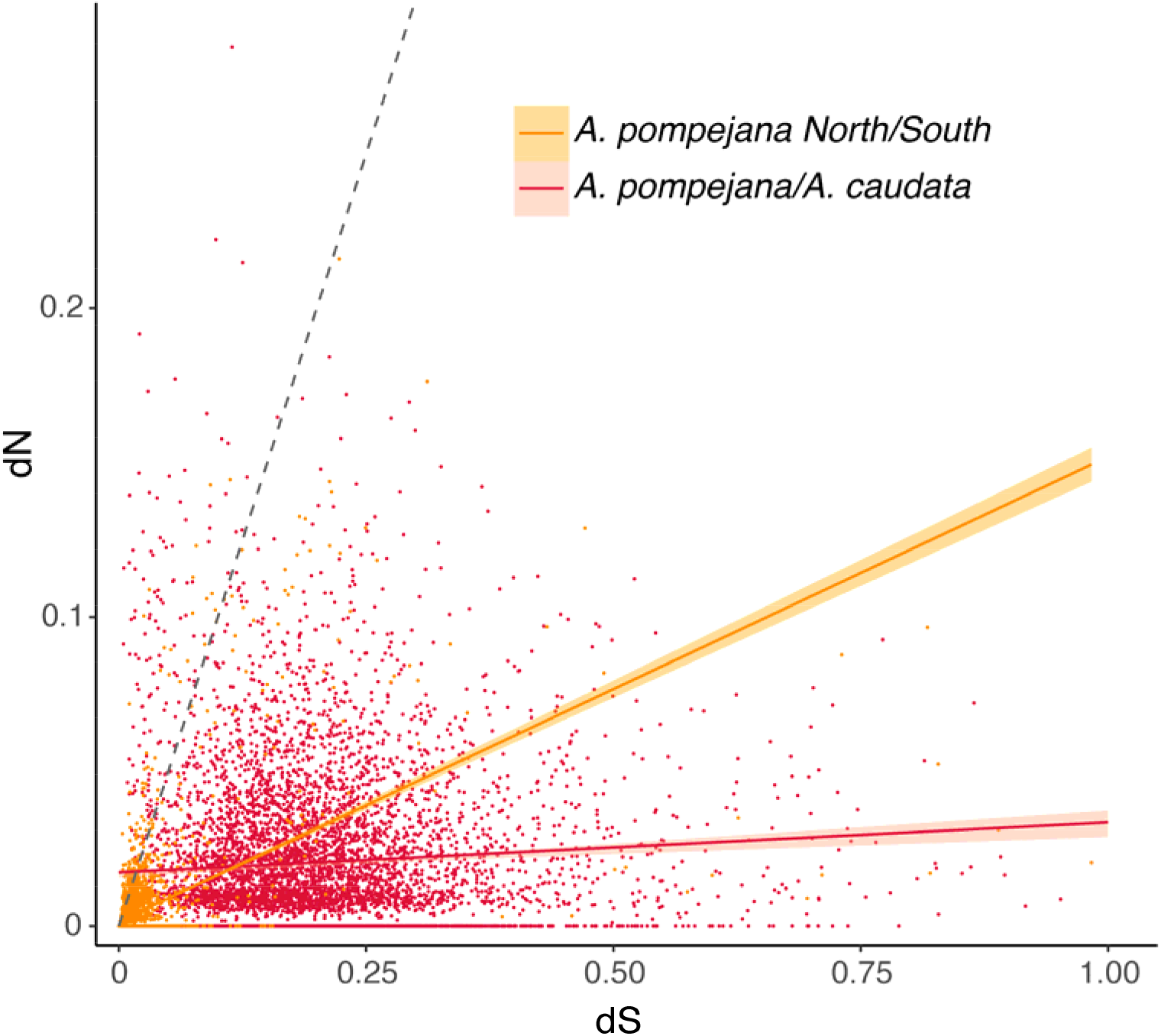
Rates of synonymous and non-synonymous divergences between *A. pompejana* North and South populations (yellow) and *A. pompejana* and *A. caudata* (red). The grey dashed line represents the expected linear relationship between d_S_ and d_N_ under neutral evolution.

The number of highly divergent genes (i.e. in the tail distribution of the genes) is almost five times greater (92 vs 20 annotated genes) and much more diversified when comparing the late and early stages of speciation, respectively. Interestingly, most genes with saturated d_S_ rates in the early step of speciation are either histone deacetylases or belong to the tubulin multigene family (Supplementary Table 1). Most α- and β-tubulins are under strong purifying selection (indicated by the relatively low rate of non-synonymous mutations) but are likely to contain at least one or two non-synonymous substitutions in their divergence, suggesting that the family may have endured a burst of duplications, and subsequent independent lineages sorting from both sides of the East Pacific Rise barrier. In contrast, most of the 92 annotated genes with a high evolutionary rate in the late step of speciation encode for a wide range of functions (e.g., replication, translation, intracellular transport…), which are quite similar to those also depicted for positively-selected genes in the Ap/Ac comparison. More specifically, highly divergent genes which also exhibited fixed non-synonymous substitutions were also found in steroid/phospholipid metabolism (phospholipid-transporting ATPase, sphingolipid delta(4)-desaturase, fatty acid-binding protein, inositol polyphosphate 5-phosphatase), carbohydrate catabolism and the tricarboxilic cycle implied in the worm’s epibiosis with specific production of acetyl-CoA (carbohydrate sulfotransferase4 dihydrolipoyllysine-residue acetyltransferase, acyl-CoA synthetase), cell motility, spermatogenesis, and egg-sperm fusion (dynein, integrin, disintegrin, protocadherin), embryonic and neuron development (methenyltetrahydrofolate synthase, protein sidekick), membrane (nidogen-1, lipoprotein receptor1, EH domain-containing protein3, transmembrane and coiled-coil domain-containing protein1, spectrin) and tegument organization (HHIP-like hedgehog protein1, biotin, α-actinin) but also, proteins more targeted on putative “adaptive” molecular pathways such as oxidative and thermal stress response (peroxidase, peroxiredoxin 4, glutathione peroxidase, Hsp70, Hsp83), DNA repairs and methylation (ATP-dependent RNA helicase, DNA ligase1, RNA polymerase II transcription subunit, adenosylhomocysteinase B, BRCA1-associated RING domain protein1, histone deacetylase3), mRNA A-to-I editing (Double-stranded RNA-specific adenosine deaminase, xanthine dehydrogenase), protein glycosylation (Protein O-linked-mannose beta-1.2-N-acetylglucosaminyltransferase1), adaptation to hypoxia (Hypoxia-inducible factor1, carbonic anhydrase) and proteins involved in protein homo-heterodimerization (complement factor H-related protein2, BTB/POZ domain-containing protein7).

### Distribution of divergence and associated d_N_/d_S_ along the genome

The distribution of gene divergence was examined over the 79 scaffolds exceeding 300,000 bp with the *A. pompejana* North/South comparison (Figure 6D). Because the global level of divergence is low and close to zero, values of divergence are homogeneously distributed with a few punctual observable outliers. As a consequence, about 83% of the examined scaffolds have no divergent genes (Figure 6A and 6D). Scaffolds containing only one divergent gene (Figure 6B and 6D) account for 15% of the examined portion of the genome and only one of the largest scaffolds displays an island of divergence pattern, and may be linked to sexual chromosome (Figure 6C and 6D). On the contrary, for the *A.caudata/A.pompejana* comparison, divergence is clearly structured along the genome in large islands of highly divergent genes for 97% of the scaffolds (Figure 6D). Each of the 76 scaffolds only carries very few genes with no divergence and we were not able to identify any region or island of no divergence among the examined scaffolds. Here, divergence is clearly structured in large blocks where all the genes and their putative exons are divergent. The average values of d_N_/d_S_ per scaffold are globally homogeneous between scaffolds but the dispersion index of the d_N_/d_S_ values around this mean fluctuates substantially. Gene density among scaffold however greatly varies and therefore impacts the number of genes involved in the genomes congealing.

**Figure 6:**
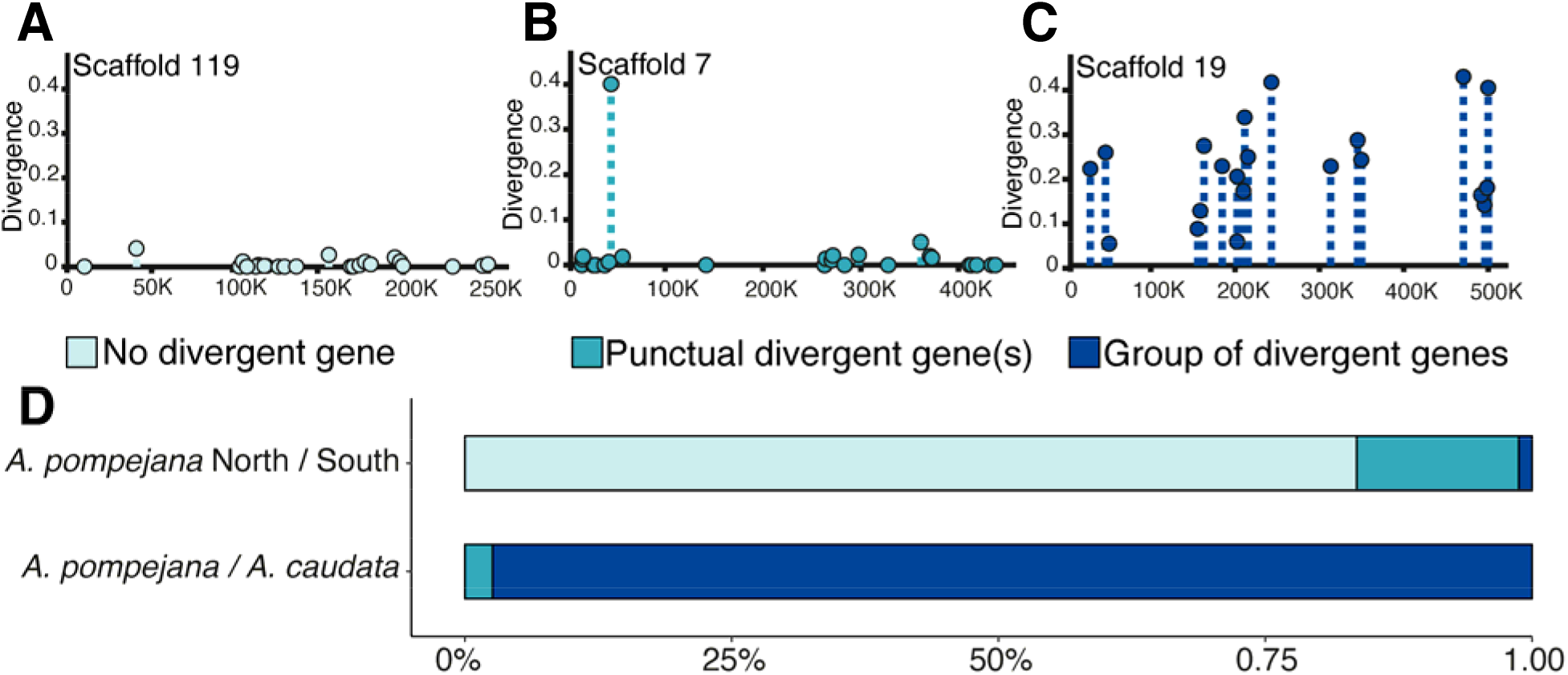
Genomic distribution of the divergence along the longest scaffolds (>400,000bp) *A. pompejana* draft genome (subset of genes representing about 1/10 of the genome). According to the index of distribution and our criterion of selection by eye, three distribution patterns of the divergence are found along scaffolds: (A) no divergent gene identified on the scaffold (in light blue), (B) sporadic presence of one divergent gene on a scaffold (in turquoise), and (C) group of spatially linked divergent genes (in dark blue). The relative proportion of each genomic pattern is shown in (D).

For both the early and late stages of speciation, no correlation between d_S_ and d_N_ has been found over the whole set of orthologous genes (The North/South *A. pompejana* comparison: r=0.18, p < 2.2e^-16^, The *A. caudata*/*A. pompejana* comparison: r=0.07, p =1.794e^-08^). We therefore estimated the Pearson’s correlation coefficient within each scaffold to uncover putative islands of divergence and analyzed the distribution of this coefficient among scaffolds. The distribution of the Pearson’s coefficient is almost bimodal and symmetrical around zero during the early stage of speciation (Figure 7A). Because most genes are weakly divergent (i.e., close to zero), this may indicate that scaffolds segregate into two distinct patterns: polymorphic genes for which allelic lineage sorting is incomplete and where the d_N_/d_S_ is biased by excesses of deleterious/artifactual mutations and (2) genes for which allelic lineage sorting is completed with a fixed divergence where positive selection may have acted. In contrast, the distribution of the correlation coefficients displayed a clear shift towards a positive correlation between divergence and associated d_N_/d_S_ values during the late stage of speciation (Figure 7B). Because most genes are highly divergent, this overall increase of d_N_/d_S_ with divergence therefore suggests that most of the genes have evolved to adapt to a new ecological niche after the separation of the two sister species.

**Figure 7:**
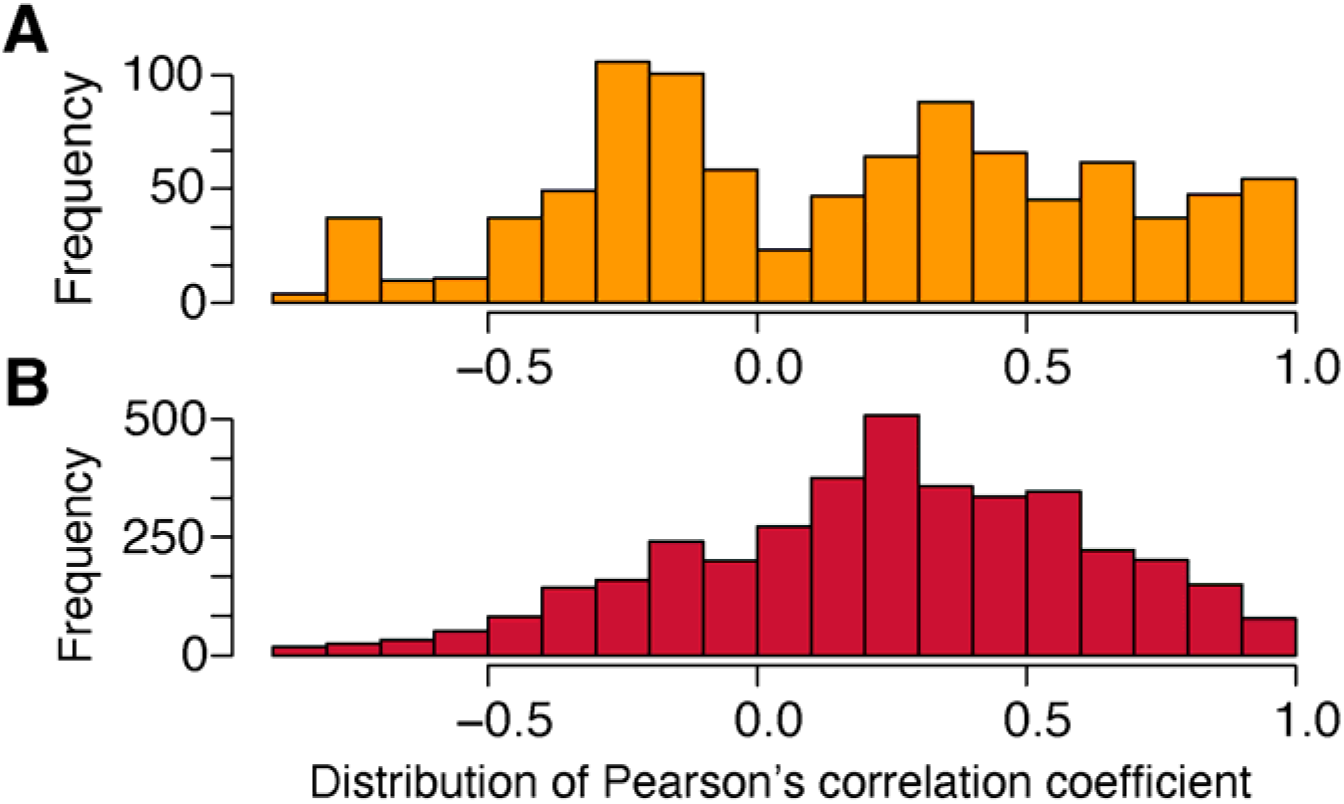
Distribution of the Pearson’s correlation coefficient between divergence and associated dN/dS values calculated for each scaffold containing more than 5 genes: (A) *A. pompejana* North/South in yellow and (B) *A. pompejana / A. caudata* in red.

## Discussion

Considering the speciation continuum from one end to the other is essential to shed light on the mechanisms by which genomes separate, and more specifically the timing of reproductive isolation and species specialization. In this study, we first examined an early stage of allopatric speciation between populations of the Pompeii worm, *Alvinella pompejana* geographically separated by a physical barrier to gene flow at the Equatorial triple junction of the East Pacific Rise and the Galapagos rift [3, 17]. This geographic separation of the Pompeii worm populations is also associated with different venting dynamics related to different spreading rates of the ridge segments that may play a crucial role in the proportion of still-hot chimneys and colder vent habitats and thus the distribution of the microbiota/biota [8, 56]. Then we also examined the molecular evolution signature associated with a much more advanced stage of the speciation process that separated *A. pompejana* and its syntopic sister species *A. caudata*, which exhibits a slightly different niche. This event is relatively old and possibly predates the separation of the EPR and the North East Pacific ridge (JdF) communities by vicariance that led for example to the separation of sibling species like *Paralvinella pandorae pandorae* and *P. p. irlandei*, about 28 Mya [1]. Mitochondrial and nuclear divergences estimated between *A. pompejana* and *A. caudata* are indeed greater than those estimated for these latter species (supplementary data in [57]).

To investigate the effect of “neutral” evolution and both “negative” and “positive” selection along genomes of separating species, we estimated both divergence and d_N_/d_S_ values from a collection of orthologous genes by comparing transcriptomes of *A. pompejana* and *A. caudata* against a first draft of *A. pompejana* genome. Although the d_N_/d_S_ ratio is determined by the combined effects of neutral, advantageous and deleterious mutations, it can tell us a lot about the general impact of natural selection on the evolution of the coding portion of the genome. In this context, a divergence scan on both synonymous and non-synonymous sites represents a very powerful tool for tracing back the role of natural selection in the semi-permeability of diverged genomes and the time at which these genomes started to congeal. It should be however noted that d_N_/d_S_ provides reliable information only for genes with complete lineage sorting where a substantial amount of substitutions have become fixed between the separated genetic entities, and caution must be taken during the early steps of speciation where most of loci are still polymorphic. On the other side of the speciation spectrum, highly divergent genomes may also contain a large number of saturated sites that leads to an overestimation of the d_N_/d_S_ ratio in fast-evolving genes.

During the early steps of speciation, the large majority of genes (about 70%) are stacked around null values of d_N_ and d_S_ but the relationship between the two rates for the remaining genes is on average 5:1, one non synonymous substitution for five synonymous ones, probably due to unfixed mutations in the worm’s polymorphism. Allele divergence and associated d_N_/d_S_ distributions between (North vs South individuals) and within (North individuals) the Pompeii worm’s populations are indeed quite similar although a greater number of higher values has been observed in the first comparison. This can be partially explained by the fact that the Equatorial barrier to gene flow is not completely sealed [15] allowing rare episodes of allele introgression. On the opposite, the slope of the regression line between d_N_ and d_S_ is close to zero during the late steps of speciation and clearly highlights a large accumulation of synonymous substitutions close to saturation for a large number of genes. This strongly suggests that even if a great number of non-synonymous substitutions are positively-selected, most of this signal is likely to have been erased in fast-evolving genes. Outlier genes associated with high evolutionary rate (d_S_>0.5) should thus represent fast-evolving genes that may have endured duplications, selective losses or gains of duplicates according to habitat/geographic isolation during the early steps of speciation, but also genes that may have already lost their ‘adaptive’ signal in the face of saturation at synonymous sites during the late steps of speciation.

As a general observation across many taxa, the average value of d_N_/d_S_ over genes between closely-related species is around 0.20 [58–60]. Multiple species comparisons suggest that more than 70% of the whole set of mutations are strongly deleterious in most species and up to nearly 30% are slightly deleterious or neutral [61]. Information gathered with the present study is consistent with nearly 87% of mutations being deleterious or slightly deleterious in *Alvinella* worms (d_N_/d_S_ mean = 0.10-0.15). This suggests that the majority of *Alvinella* genes are under strong purifying selection probably as a consequence of the extreme thermal conditions encountered by the worms [57]. Given such background selection, it is highly possible that many proteins with d_N_/d_S_ ranging from 0.5 to 1 share some history of positive selection, without any possibility to test it.

### Early steps of speciation: lack of genes implicated in reproductive isolation despite accumulation of divergence at many genes?

Results from our genome-wide study are consistent with previous results relying on the mitochondrial genome and some other genes in showing the geographic isolation of *A. pompejana* populations across the Equator [15,62–63]. We indeed observed a clear signal of divergence at nearly 30% of genes between the transcriptomes of the southern and northern individuals of *A. pompejana*, despite a huge number of non-divergent genes (55% of the total number of genes examined). This emerging divergence corresponds to a modal value of 0.025 substitution per codon and represents a 4-times increase of both transitional and transversional substitutions when compared with pairwise alignments of the same orthologous genes between two distinct northern individuals from the same population. This emerging divergence at the genome scale confirms the ongoing allopatric speciation occurring for these lineages along the EPR possibly reinforced by local adaptation. Time since the population splitting does not seem to have been however sufficient to produce a specific genomic architecture of divergence (*i.e.,* islands of speciation) as most of the divergent genes are scattered among scaffolds. This may be explained by the fact that the predicted North-to-South migration, though very low [15] could still be able to overcome population differentiation over large portions of the genome and thus, would prevent genes from accumulating fixed divergence. Alternatively, time since population splitting may still be too short to result in a complete lineage sorting for most genes and the subsequent emergence of genetic incompatibilities. According to Orr & Turelli [64] the strength of reproductive isolation (i.e. fitness load of putative hybrids) increases as a function of the squared number of newly fixed mutations in separated populations. The number of fixed substitutions (and hence associated genetic incompatibilities) between the two isolated entities represents 15560 substitutions in coding sequences (as the product of 2,075 300-codons genes with a divergence of 0.025 substitutions per codon (30% of the investigated genes)) for approximately 1/10 of the whole genome. Compared to other eukaryotic models (*e.g.*, about 70,000 changes accumulated between sub-species of the *Drosophila simulans* complex over 0.25 My [65] for about the same number of coding sequences), the number of accumulated changes (i.e. ∼15,560) is quite low if the two populations separated 1.5 Mya [15], even if the generation time may be about 10 times greater in *Drosophila*. Such discrepancy might be explained by a strong stabilizing selection associated with these extreme but constant environmental conditions over space and time. Gene networks models indeed predict that the hybrid incompatibility dynamics may be greatly reduced by stabilizing selection while producing a “basin of attraction” for the optimal genotype on both sides of the barrier to gene flow [66].

In this early stage of speciation, very few genes showed evidence of positive selection and these genes do not relate to genes currently depicted as involved in reproductive isolation (*e.g.*, genes involved in gamete recognition, pheromones, mito-nuclear incompatibilities, hybrid sterility or immune incompatibility [67–70]. Neither sperm-egg binding or egg-fusion proteins nor nuclear proteins transferred to the mitochondria displayed signatures of positive selection or strong divergence. However, almost all genes showing high evolution rates in our data encode for tubulins, which represents highly diversified multigene families and can have a great influence on cell division (mitosis/meiosis) and the spermatozoon flagellum architecture (which is extremely reduced as a byproduct of the internal fertilization of oocytes in *Alvinella* [71]. It should also be noted that a meaningful proportion of positively-selected genes could indirectly participate to the establishment of reproductive barriers as they encode for proteins involved in spermatogenesis, the development of gonoducts, the sensory system and setae/hooks that may play a role in male and female pairings (*i.e.*, pre-zygotic isolation). Other genes under divergent selection are mostly unknown or involved in immunity and endocytosis and may affect the interaction of the worm with the associated campylobacteria-dominated microbial assemblages covering its tube and its dorsal epidermis (*i.e.*, trophic and detoxification role of the worm’s epibiosis [72]). Microbial communities are likely to be genetically different between vent fields as previously observed for other vent taxa [73] and thus could induce co-evolutionary pathways of divergence in the worm’s immune system [74]. In the specific cases of sympatric or parapatric speciation, genes under positive selection tend to be predominant among rapidly evolving genes [75]. Synergic effects between positively-selected genes (linkage disequilibrium) are likely to produce local barriers to gene flow if their number is high enough [30, 46], but this is obviously not the case here. Analysis of the correlation between divergence and d_N_/d_S_ revealed a potential antagonistic action of selection on divergent genes, one half of scaffolds producing a negative correlation between the two parameters whereas the other half exhibited a positive correlation. Such correlations are clearly biased by the very high number of genes with no divergence but negative correlations over many scaffolds could be interpreted as an excess of deleterious mutations due to incompletely sorted polymorphisms for many genes. Additional work on the genetic differentiation of the two separating lineages is currently undergone at both the genome and population scales using ddRAD markers to complement this present study.

### Late stage of speciation: role of gene specialization and gamete recognition genes in species reinforcement after genome congealing

As opposed to the recent history of allopatric speciation in *A. pompejana*, the scan of gene divergence (*t*, d_N_, d_S_) between *A. pompejana* and its syntopic sister species *A. caudata* provides a contrasting story of gene specialization after the congealing phase of the two corresponding genomes. Our results confirm previous studies [51–52], who showed that these species did not share any allele at enzyme loci and refuted the initial hypothesis that the two worms corresponded to two morphological ontogenetic forms of the same species [49]. Both species occupy the same hydrothermal vent habitat (hottest part of vent chimneys) and have exactly the same geographic range from 21°N/EPR to 38°S/PAR (Pacific-Antarctic Ridge) leading to the possibility of sympatric speciation. The two species indeed live in a spatially heterogeneous environment, which greatly varies in time [76–77]. According to Gaill et al. [78], the parchment-like tube of the two worms is highly thermostable, and the two species share many adaptations allowing them to thrive in these waters loaded with heavy metals and sulfidic compounds (*e.g.*, similar epibiotic flora composition: [79], similar lipids in the cuticle: [80], similar branchial crown, gaz transfer structure and haemoglobins [81–82], and same reproductive mode: [71]). Our study of gene divergence however reveals a very old separation of the two species and a relatively high proportion of genes (2%) still showing evidence of positive selection when considering that the average gene divergence is very high (*i.e.*, close to 0.2 substitution per synonymous site). Most of the positively-selected genes are not necessarily those with the highest rate of evolution, thus indicating that the signal of positive selection may have been erased by excesses of synonymous substitutions for many genes. Distribution of gene divergence appears quite homogeneous among scaffolds in terms of basal divergence, which is a strong indication that a genome-wide congealing effect occurred much earlier. We estimated the time of genome congealing around 26.5 Mya by using the substitution rate previously estimated by Chevaldonné *et al.* [83]. The first mapping of divergence and associated d_N_/d_S_ showed however a strong heterogeneity of these variables along the genome with some scaffolds having relatively higher divergence values. Although we cannot rule out that these scaffolds could represent either regions of lower recombination rates or a portion of sex-linked chromosome, their high divergence could also trace back to the early steps of the speciation process (i.e. primary islands of speciation), in this case suggesting that the separation of the two species may have been initiated much more earlier.

Gene divergence is also slightly positively correlated with d_N_/d_S_ within each scaffold. This therefore suggests that the increase in gene divergence have been accompanied by the accumulation of non-synonymous mutations. Assuming that d_N_/d_S_ only increase when synonymous sites become saturated, this trend could either indicate an adaptive divergence associated with a long period of gene specialization for the two sister species regarding their habitat after their separation, or that the time since species separation is so high that most genes reached saturation but this later interpretation does not fit well with the presence of positively-selected genes. The genes under positive selection (d_N_/d_S_ > 1) are more specifically involved in the repair/replication/biosynthesis of nucleic acids but also in several metabolic pathways usually involved in the stress response, carbohydrate catabolism and lipid/steroid metabolisms associated with membrane organization and tegument formation. Targets of positive selection seem to be consistent with gene specialization in response to different thermo-chemical regimes encountered by the worms, and more specifically high temperatures, natural doses of radioactivity and high concentrations of sulfides and heavy metals producing reactive oxygen species (ROS) and reactive sulfur species (RSS), and suggest that the two species do not share exactly the same niche. Annotation of fast evolving genes also revealed that most of them belonged to exactly the same metabolic pathways impacted by positive selection including membrane/tegument organization. Regarding fast evolving genes, we noted a net enrichment in proteins involved in oxidative stress response (peroxidase, Hsps), adaptation to hypoxia, DNA repairs and methylation, mRNA A-to-I editing, and protein glycolysation. The two latter molecular processes represent a powerful way to adapt to differing thermal regimes [84–85]. These results therefore indicate that the adaptive evolution of the two species has been a sufficiently ‘old’ process to erase most signatures of positive selection that has likely played a crucial role in their ecological isolation. As opposed to *A. caudata*, *A. pompejana* is one of the first colonizer of newly-formed and still hot chimneys [86] when these edifices are made of porous anhydrite (barium/calcium sulfates) with temperatures above 100°C. Consistently, slight differences in terms of protein composition with more charged residues in *A. pompejana* and more hydrophobic/aromatic ones in *A. caudata* have been described [57]. Interestingly, when looking at the proportion of each species between chimneys at a local spatial scale (*i.e.*, hundreds of meters), it can be observed that newly-formed anhydrite edifices and ‘white smokers’ are mostly inhabited by *A. pompejana*, whereas older edifices and ‘black smokers’ are mainly characterized by a greater abundance of *A. caudata* [7]. Accelerated differentiation of genes involved in the adaptive response of the worms to environmental variations are likely to explain their different ecologies and behaviors to cope with ‘high’ temperatures. For instance, tubes of the two species are closely entangled in colonies but with differences in their opening [72] (D. Jollivet, *pers. obs*.). While the aperture of the tube of *A. pompejana* is widely opened like a funnel with septa, that of *A. caudata* is ended by a small hole. The shape of the tubes is likely related to the thermoregulatory behavior of the worms that helps individuals to refresh their microenvironment through water pumping [77, 87]. While *A. pompejana* is widely observed moving inside and outside its tube to refresh it with the cold surrounding waters, *A. caudata* usually stays in its tube but exhibits a tail without organs used to cultivate its epibiotic flora that may be viewed as a thermal sensor to position the worm at exactly the right temperature.

Despite this very long history of speciation and subsequent species specialization, one of the most striking observations we made is that many genes involved in gamete recognition (*e.g.*, zonadhesin, innexin) are under positive selection, suggesting that species sympatry may still play a role in species reinforcement. These positively-selected proteins are accompanied by several fast-evolving proteins involved in spermatogenesis, and egg-sperm fusion (*e.g.*, dynein, integrin, disintegrin, protocadherin), and embryonic development (*e.g.*, methenyltetrahydrofolate synthase). Moreover, specific pathways such as the Acetyl-CoA cycle (3 targets) together with a lipid desaturase seems to become highly differentiated. In *Drosophila*, desaturase genes and the Acetyl-CoA cycle are both involved in the production of cuticular hydrocarbons (CHCs) which have a crucial role in adaptation to desiccation and also act as pheromones involved in mating behavior [70]. It is therefore possible that these two worm species preliminary evolved in a different geographical context for a long period of time and then met secondarily during the recolonization of the EPR after the episodes of tectonic isolation that structured the hydrothermal fauna between 11-18 Mya as recently suggested to explain the phylogeographic patterns of the *Lepetodrilus elevatus* complex of limpet species [4]. This rather late evolution of prezygotic mechanisms of isolation contrasts with the allopatric situation encountered between the northern and southern populations of *A. pompejana* and supposes that these mechanisms could last for a very long time even after a genetic barrier has been erected.

## Conclusion

Our analyses pointed out the non-negligible role of natural selection on both the early and late stages of speciation in the emblematic thermophilic worms living on the walls of deep-sea hydrothermal chimneys. They shed ligth on the evolution of gene divergence during the process of speciation and species specialization over time. Due to habitat fragmentation, populations separate and start to accumulate putative adaptive mutations (and possibly genetic incompatibilities) in allopatry where they find differing local conditions but are likely to consolidate reproductive isolation in sympatry following secondary contacts during the specialization process, even after genomes diverged almost completely (congealing effet). These first analyses raise many further questions about the evolutionary mechanisms that led to the speciation of *Alvinella* spp. and their subsequent distribution within the spatial micro-mosaic of habitats typifying hydrothermal vents. Additional studies combining polymorphism and divergence are needed to better understand the respective roles of geographic and ecological histories of the worm’s populations in speciation in a such fragmented and instable environment.

## Material and methods

### 1) Animal sampling and genome/transcriptome sequencing

The genome of the Pompeii worm was assembled from a single individual collected at 9°50N/EPR. Several DNA purification methods using alcohol precipitation and/or column purification resulted in DNA samples that could not be efffectively digested by restriction enzymes. To reduce mucopolysaccharides or similar co-precipitating contaminants, we purified the DNA from worm tissue with phenol extraction and isopropanol precipitation followed by a clean-up step with cesium chloride gradient ultracentrifugation [88]. The DNA was used to prepare paired-end shotgun and 5 kb mate-pair libraries for Illumina genome sequencing [89].

We used at least two additional individuals of *A. pompejana* and *A. caudata* coming from the same locality to perform RNA sequencing and the subsequent assembly of transcriptomes for the northern specimens of the two species [57]. In parallel, a Sanger sequencing project of a reference and fully annotated transcriptome was performed using several individuals coming from a single chimney of the hydrothermal vent field 18°25S [90], for additional information about this annotated transcriptome) and, represented the third transcriptome used to characterize the specimens of *A. pompejana* located further south to the Equatorial barrier to gene flow previously described [3, 15].

### 2) Sequencing (see Metrics in Supplementary Table 2)

- Genome and transcriptomes

Genome of the Pompeii worm (about 370 Mb) was sequenced from a single individual using both a shotgun and mate-pair 5 kb sequencing using the Illumina HiSeq2500 technology at the A*Star Molecular and Cell Biology Institute (*Alvinella* genome project: coord. A. Claridge-Chang). Paired-end reads libraries (insert size 300bp) were assembled using SOAPdenovo2 (v2.0, -K 41 -d -R), contigs were then scaffolded with OPERA (v1.3.1) using both paired-end (insert size 300bp) and mate pair libraries (insert size 5 kb). Finally, the assembly was gapfilled with Gapcloser (v1.12) to subsequently produce about 3000 scaffolds with a size greater than 15 kb.

The annotated transcriptome of the southern form of the Pompeii worm collected at the vent field 18°25S has been already published in 2010 [90], and transcriptomes of the northern forms of *A. pompejana* and *A. caudata* have been already published in 2017 [57] (see Availability of data and materials). All transcriptomes were assembled *de novo* using the software Trinity [91], the sequencing effort was a quarter of a lane (40million reads) per species on a HiSeq 2000 at Genome Québec.

### 3) Estimation of divergence and d_N_/d_S_ for pairs of orthologous genes

We designed a customized pipeline (described below) in order to calculate divergence and estimate the selective pressures along the coding regions of assembled transcriptomes of closely related species previously positioned on our reference genome, using a 24-columns MegaBLAST outputs. The code to perform analyses for this study is available as a git-based version control repository on GitHub (https://github.com/CamilleTB/dNdS_cartography_Alvinella_speciation)

- Megablast

Because coding sequences are compared between divergent individuals within a given species or between closely-related species, we performed a sequence-similarity search using each collection of transcripts obtained for *A. caudata* and the two geographic forms of *A. pompejana* against the entire *Alvinella pompejana’s* genome with the software MegaBLAST implemented in NCBI BLAST+ 2.2.30 nucleotide-nucleotide search, optimized for very similar sequences (set expectation value cutoff = 10^-15^, maximum hits to show = 1).

- Paralog

The 24-columns MegaBLAST output was then filtered using a custom-made Perl script (called “Paralog”) to discriminate genes according to their position and occurrences in the genome. This allowed us to rule out putative duplicated genes in our subsequent analyses. Based on the positions of the hits on both scaffolds and transcripts, genes were separated into distinct categories, namely scaffold paralog, tandemly repeated paralogs, allelic forms of a single gene in either genes with or without introns. Scaffold paralogs correspond to coding sequences found on distinct scaffolds, tandemly-repeated genes to sequences found at non-overlapping positions onto the same scaffold, allelic forms of a gene corresponded to several transcripts matching exactly the same positions onto a scaffold, and the two others categories were orthologous transcripts only found once into the genome at a given scaffold position with a discrimination exon/uniq when the Megablast hits corresponded or not to a succession of fragments with different positions both onto the scaffold and the transcript (i.e. exonic regions). Paired orthologous sequences obtained in the two last categories (exon and uniq) were then used to estimate divergence and associated d_N_/d_S_ ratios in the pipeline.

- Search for coding DNA sequences (CDS) in pairwise alignments

Nucleic acid sequences were translated in the six reading frames to select for the longest CDS between two subsequent stop codons. Coding sequences containing gaps, undetermined nucleic acids (N) or alternative frames without stop were removed. To avoid false positives and reduce the risk of calculating erroneous d_N_/d_S_ values from non-coding or incorrectly framed sequences, we performed a sequence-similarity search against the UniProt database using BlastP from NCBI BLAST+ 2.2.30 (set expectation value cutoff = 10^-10^, maximum hits to show = 1). Sequences with no annotation from UniProt with a CDS shorter than 300 nucleotides were removed from the analysis.

- Divergence and d_N_/d_S_ calculation

Estimation of divergence, synonymous and nonsynonymous substitution rates and detection of positive selection in protein-coding DNA sequences were performed using the program yn00 from pamlX 1.2 in batch [92–93]. This program implements the method of Yang & Nielsen (implemented in [92]) to calculate d_S_ and d_N_ values from pairwise comparisons of protein-coding DNA sequences. The weighing parameter that decides whether equal or unequal weighing will be used when counting differences between codons was set to zero (= equal pathways between codons) and we used the universal genetic code (icode = 0). Output from yn00 was then parsed and subsequently filtered from infinite values (99) when S.d_S_ was equal to zero. When both d_N_ and d_S_ values were null (about 70% of values obtained when comparing the northern and southern forms of *A. pompejana*), d_N_/d_S_ was reset to zero. To avoid a division by zero, if d_N_ was different from zero but d_S_ was zero, S.d_S_ was reset to 1 assuming that at least one synonymous mutation was missed by chance in the sequence.

To avoid any potential effect of greater divergence at non-synonymous sites possibly due to sequencing errors when values of neutral divergence (d_S_) departed from zero, the significance of all d_N_/d_S_ values greater than zero was tested using a custom-made pipeline. For that purpose, we produced 1,000 random alignments of each pair of coding sequences using a bootstrap resampling of codons (from the bootstrap option of codeML in PamlX 1.3.1). We then estimated the associated d_N_ and d_S_ for each resampled alignment, and calculated the difference (D) between each d_N_ and d_S_ resampled values in order to test whether the observed value of this difference was significantly greater than zero (*i.e.*, fall outside the distribution of the pseudo-replicates). To this end, we performed a unidirectional one-sample *z*-test for each resampled set of paired sequences and adjusted p-values over the whole set of genes using the false discovery rate correction representing the expected proportion of false positives [94].

### 4) Identification of islands of divergence and/or adaptation

To investigate the genomic architecture of molecular divergence of putative genes under strong purifying (d_N_/d_S_ close to zero) or positive selection (d_N_/d_S_ >1) along chromosomes, both the divergence and their d_N_/d_S_ associated values were positioned along the longest scaffolds (>300,000 bp with more than 5 genes per scaffold) of the draft genome of *A. pompejana* for each pairwise comparisons (*i.e.*, AP north vs AP south, and AP vs AC). According to the gene distribution along each scaffold, three different types of patterns were characterized depending on the number and level of clustering of divergent genes (divergence > 0.1) among scaffolds: (1) scaffolds without divergent genes, (2) scaffolds with one or two isolated divergent genes and, (3) scaffolds carrying a genes cluster of divergence corresponding to a putative island of speciation. The proportion of each type was then calculated for the two sets of analyses. In addition, we calculated the mean scaffold distance (d) between two divergent genes and its standard deviation (s^2^(d)) within each scaffold. We then tested for contiguous distribution patterns of divergence along each scaffold thanks to the Fisher’s index of dispersion (s^2^(d)/mean(d)). Divergent genes are randomly distributed when the index is equal to one, over-dispersed for values lower than one and aggregated for values greater than one.

We finally tested for a positive correlation between d_N_/d_S_ and divergence (t) along scaffolds (>300,000 bp with more than 5 genes per scaffold) using a home-made R script. Such a correlation should indicate a genome-wide adaptive evolution of proteins assuming that, under neutral evolution, d_N_/d_S_ remains relatively constant before the time at which most of the synonymous sites become saturated, and then slightly decreases with longer time of separate evolution.

## Supporting information

Supplementary table 1 - genes annotation

Supplementary table 2 - genome and transcriptomes metrics

## Abbreviations

AP: Alvinella pompejana
AC: Alvinella caudata
EPR: East Pacific Rise
JDF: Juan de Fuca Ridge
FDR: False discovery rate
MYA: Million years ago

## Acknowlegments

We would like to thank the captain and crews of the research vessel L’Atalante and the pilots and associated teams of the manned submersible Nautile without which no specimen would have been sampled. We are also very grateful to the chief-scientists of the oceanographic cruises Biospeedo 2004 and Mescal 2010 for collecting *Alvinella* specimens and to S.C. Cary who kindly provided in 2006 the AP specimen used to produce the genome. We thank Romain P. Boisseau for considerable feedback on the manuscript and Ulysse Guyet for useful discussions on the statistical analyses.

## Authors’ contributions

CTB and DJ performed analyses and wrote the manuscript. EB and DJ conceptualized, designed and co-supervised the study. SH and DJ were in charge of the funded research project. ACC supervised the AP genome project. DB, NN, RC and EC were involved in the AP genome assembly, transcriptome assembly and mapping, and gene annotation. DJ, SH, and ACC edited and contributed to the proofreading of the manuscript. All authors read and approved the final manuscript.

## Availability of data and materials

The AP genome: Two replicate runs of Illumina were performed onto the same individual (2 sets of R1 and R2 compressed files of 33 Go: cf. SRA project under experiment accession numbers ERX5990023-25; ERX5990051 and, run accession number: ERR6358489-91; ERR6358517) and, used in combination to generate the first genome draft (V1). The reads of transcriptomes are available from the Sequence Read Archive (SRA) database under accession numbers SRP076529 (Bioproject number: PRJNA325100) and SPX055399-402 (Bioproject number: PRJNA80027)), and from the EST database under accession numbers FP489021 to FP539727 and FP539730 to FP565142.

## Declarations

### Ethics approval and consent to participate

Not applicable.

### Consent for publication

Not applicable.

### Authors details

^1^Present address: Emlen Lab - BRB108, Division of Biological Sciences, The University of Montana, Missoula, MT, USA

^2^Sorbonne Université, CNRS, UMR 7144 AD2M, Station Biologique de Roscoff, Place Georges Teissier CS90074, 29688 Roscoff, France

